# The Effect of Visual Field Occlusion on Visually Induced Motion Sickness

**DOI:** 10.1101/2024.12.26.630441

**Authors:** Takatsugu Aihara, Sachiko Yamada, Hiroshi Ban

## Abstract

Visually induced motion sickness (VIMS), characterized by symptoms such as dizziness, nausea, and postural instability, presents significant challenges in modern visual and interactive technologies. These symptoms undermine the immersion into and comfort of virtual and augmented reality (VR/AR) systems and reduce the usability of autonomous vehicles, highlighting the urgent need to address VIMS for broader adoption of these technologies. While previous studies have largely focused on the motion characteristics that induce VIMS, the impact of visual field configuration, such as segmentation and occlusion, has been largely unexplored. To bridge this gap, we conducted five human psychophysics experiments systematically investigating how visual field segmentation and masking influence VIMS under controlled three-dimensional optical flow conditions. Participants were exposed to different masking patterns, including variations in the number of divided sections, the masking of the central screen region, and the thickness of the masks. Paired-comparison methods were employed to assess participants’ susceptibility to VIMS under each condition. Our findings showed that visual field segmentation significantly influenced VIMS susceptibility. Specifically, the results revealed three key findings: (1) segmenting the visual field into multiple subsections using visual masks significantly increased VIMS susceptibility, (2) thinner masks exacerbated motion sickness more than thicker ones, and (3) masking the central region of the screen reduced susceptibility. These effects were independent of the presented motion patterns. Our findings offer new insights for optimizing the design of VR/AR systems, graphical user interfaces, and autonomous vehicles by highlighting the importance of visual field configuration in mitigating VIMS for improving user experience.

## 1. Introduction

Motion sickness (MS) is induced while watching movies, playing video games, or traveling by car, ship, or airplane. MS is characterized by the presence of unpleasant symptoms such as general feelings of discomfort, dizziness, fatigue, difficulty concentrating, drowsiness, headache, pallor, sweating, increased salivation, stomach awareness, nausea, and vomiting. Individuals with a normal vestibular system have been found to experience MS, whereas those with an impaired vestibular system do not, indicating the prominent role of the vestibular system in MS (Paillard et al., 2013). MS is not a disease but a natural and transient response to provocative motion or sensory stimuli. Therefore, MS may occur in healthy individuals depending on the motion or sensory stimuli and physical condition; however, there may be individual differences in symptoms and their severity. Infants are more resistant to MS, and susceptibility to MS starts to increase at approximately 2 years of age, peaks during adolescence, and then gradually diminishes, though not entirely, in adulthood. It has been reported that females are more susceptible to MS than males (Paillard et al., 2013). Given its impact on individuals of all ages, understanding the underlying psychological and neural mechanisms of MS is crucial for reducing discomfort in virtual reality (VR) environments, which is essential for achieving high levels of immersion and ensuring a pleasant user experience, ensuring comfortable travel experiences via a car or other modalities and developing effective countermeasures.

The sensory conflict theory is the most prevailing theory regarding MS. The sensory conflict theory posits that a discrepancy among or within visual, vestibular, and somatosensory information leads to the incidence of MS (for the refined version, please refer to the neural mismatch model by Reason (1978)). It is well-documented that rear-seat passengers, especially children, in automobiles or buses are more susceptible to experiencing MS (Gordon & Shupak, 1999; Perrin et al., 2013) owing to the incongruity between the visual and vestibular information caused by obstruction of the view outside the vehicle by the front-seats and front-seat passengers. Individuals may experience MS-like symptoms when exposed to visual motion without corresponding body movements; this phenomenon is referred to as visually-induced motion sickness (VIMS). The prevalence of VIMS has increased in recent years, which is attributed to the recent advancements in video-game technologies, online video-sharing systems, television, and movies. In addition, powerful, realistic, and captivating visual contents displayed using large immersive screens, three-dimensional (3D) stereoscopic video displays, and VR systems may render the audiences more susceptible to VIMS if the display properties and parameters are not adequately adjusted. The VIMS resulting from exposure to VR environments is also called cybersickness. Extensive exposure to such provocative visual motion stimuli in daily life has contributed to an increase in the incidence of VIMS in recent years. In addition, the use of automated vehicles has been facilitated by recent technological innovations and the expeditious enactment of pertinent laws and regulations. It is predicted that vehicle automation will amplify the incidence and intensity of MS (Diels & Bos, 2015).

Thus, the implementation of countermeasures is imperative for the mitigation of MS, thereby facilitating the enjoyment of visual contents and the use of automated vehicles. Two distinct types of countermeasures have been discerned: the first measure focuses on the viewer’s perspective, whereas the second measure entails the presentation of visual stimuli. Repeated exposure to stimuli that can induce VIMS may lead to symptom alleviation through adaptation. However, this approach lacks practical feasibility. A more pragmatic strategy involves the use of pharmacological agents to alleviate symptoms of MS, such as centrally acting anticholinergics, antihistamines with antimuscarinic activity, and combinations of anticholinergic and adrenergic agents (Gordon & Shupak, 1999). Scopolamine, an anticholinergic, is the most effective drug for the management of MS. However, it can cause significant side effects, such as dry mouth, drowsiness, blurred vision, and confusion. In addition, the prolonged use of transdermal scopolamine increases the risk of adverse effects and may lead to drug dependence. Antihistamines are the most administered drugs for the management of MS in pediatric cases. However, these agents are associated with the incidence of adverse effects, such as drowsiness and dry mouth, although they are less severe than those observed with the use of oral and parenteral scopolamine. Thus, it may be useful to develop an alternative strategy to alleviate MS by optimizing the method of visual presentation. Therefore, in this study, we examined how the screen design, such as occlusion, subdivision of the display, can modulate an observer’s impression of MS.

The effect of occlusion has been studied in the field of screen design independent of its interaction with VIMS. Amodal completion, a well-known phenomenon related to visual occlusion, is the apparent ability of the visual system to visualize an entire object by filling in the missing or occluded sections (Gerbino, 2020). For instance, the moon is perceived as a holistic circular form even when it is partially occluded behind a building. Various visual illusions related to object occlusion have been reported wherein the perceived dimensions, configurations, and positions of the partially occluded objects exhibit distortions. Occlusion illusion and Poggendorff illusion must be considered when designing comfortable screens. Occlusion illusion is a visual illusion wherein the visible portion of a circle (e.g., a semicircle), which is partially occluded behind a rectangle, appears to be larger than the physically identical semicircle presented in isolation. This illusion can be explained using Emmert’s Law, which states that the perceived size increases with distance if the retinal size remains constant. Thus, if the observer perceives that the occluder (i.e., the rectangle) is in the same depth plane as the isolated semicircle, then the occluded semicircle must be farther away than the isolated semicircle, and larger (But please note that there is another explanation by Palmer et al. (2007)). The Poggendorff illusion is a visual illusion wherein collinear oblique lines appear misaligned when the middle portion of the oblique line is occluded by a rectangle or parallel lines. According to Emmert’s law again, the rectangle or the space between the vertical lines appears to shrink if it is perceived as nearer. The shrinkage drags the two oblique lines inwards; therefore, they appear misaligned. This explanation was supported by the results of a series of experiments (Talasli & Inan, 2015). These phenomena related to the visual occlusion suggest that the occluder causes a perceptual distortion of the partially visible objects. Such distortions of visual scenes may inadvertently enhance VIMS by distorting the trajectories of moving objects. Therefore, in this study, we investigated the impact of scene subdivision and occlusion on the impression of VIMS.

Previous studies have investigated the different perceptual characteristics of central and peripheral vision in VIMS and related phenomena, such as vection, and postural stability. It was found that the symptoms of MS, vection, and induced body movement reduce progressively when the peripheral visual field is occluded (Perrin et al., 2013; Kawakita et al., 2000; Kim & Kim, 2019; Piponnier et al., 2009; Horiuchi et al., 2017; Raffi & Piras, 2019). The central circular (peripheral vision) or peripheral (central vision) area was masked in these studies. To the best of our knowledge, no previous study has systematically investigated whether VIMS and its related phenomena are affected when the visual field or screen is masked or subdivided in a different manner. Investigating the effects of different scene appearances on VIMS susceptibility as well as the effect of the difference between central and peripheral vision has important implications. As a first step, the present study focused on the fact that rear-seat passengers in a car or bus are more susceptible to developing MS. Their field of vision is occluded or divided into right and left sections by the seats, headrests, and pillars, which can cause MS. Therefore, several masking patterns inspired by seats and pillars were prepared in the present study. It was hypothesized that the following aspects of visual design may affect VIMS: the number of divided displays, masking of the focus of expansion (FOE) or the screen center, and the area and width of the mask. Paired-comparison experiments were conducted to test these hypotheses.

## 2. Material and Methods

### 2.1 Overview

The participants were instructed to rate the strength of the subjective impressions of VIMS for various optical flow stimuli in which sections of the visual field were systematically occluded, and only sections of the dot motions were visible. The judgement was ranked using Thurstone’s paired comparison method (Thurstone, 1927). Five experiments were conducted utilizing different occlusion patterns and dot motion conditions. A paired comparison was performed to evaluate VIMS because the preliminary experiment indicated that the simulator sickness questionnaire (SSQ) lacked sufficient sensitivity to distinguish differences among masking patterns. Here, SSQ is typically used in motion sickness studies; however, it was inadequate in this study. This may be because the SSQ is designed to describe the current state of sickness rather than to predict an impending onset of sickness. The followings were common to all five experiments. The experiments were conducted in accordance with the Ethics Codes of the Center for Information and Neural Networks (CiNet), National Institute of Information and Communications Technology (NICT), and were approved in advance by the local ethics and safety committees at CiNet and NICT. All participants provided informed consent before enrolling in the experiment. The participants were informed that the experiment could induce mild VIMS; however, they were not informed regarding the specific purpose of the study. The participants had normal or corrected-to-normal vision, with no reported history of neurological or psychiatric disorders. A total of 187 individuals participated in the study and received cash awards in return for participating. Some participants participated in multiple experiments on different days.

### 2.2 Visual Stimuli

Visual stimuli were rendered in three-dimension (3D) by binocular disparity to enhance the realistic nature of the movie, thereby increasing the possibility of inducing VIMS symptoms. White dots (1200 dots, diameter: 0-9 pixels, CIE 1931 (x, y) = (0.3110, 0.3218), luminance = 64.90 cd/m^2^ [measured with Konica-Minolta CS150]) were displayed in random positions on a dark gray background (CIE1931 (x, y) = (0.3130, 0.3218), luminance = 5.07 cd/m^2^). The movement of the dots was different in each experiment (the movements are described in detail in each subsection). The GLSL shader technique implemented in the ‘moglDrawDots3D’ function from the Psychtoolbox-3 MATLAB library (Brainard, 1997; Pelli, 1997) was used to efficiently draw a large number of dots in 3D. The masks were drawn as black rectangles (CIE 1931 (x, y) = (0.3192, 0.3351), luminance = 1.09 cd/m^2^). Each experiment had seven masking patterns, including a no-mask condition, and these patterns were different in each experiment (the masking patterns are described in detail in each subsection and illustrated in Figures 2, 3, and 4). The movies provided as Supplementary Information present examples of visual stimuli.

### 2.3 Apparatus, tasks, and procedures

The experiments were conducted in a sound-proof booth (Avitex AMC4025; Yamaha Corporation, Japan). All stimuli were presented on a 27-inch (59.7 cm×33.6 cm) LCD monitor (ROG SWIFT PG278Q, ASUS, Taiwan; resolution 2560×1440 pixels; refresh rate 120 Hz). The participants were made to wear active shutter goggles (3D Vision2 Kit; nVidia, USA) to enable a stereoscopic 3D experience. An in-house stereo vision acuity screening tool (https://github.com/hiroshiban/StereoScreening) was used to confirm the absence of problems in stereo vision before the main experiments. The viewing distance in Experiments 1, 2, and 3 was 81.8 cm. However, the viewing distance was reduced to 40.9 cm in Experiments 4 and 5 to widen the viewing field and enhance the immersiveness of the visual stimuli. The field of view was 40.1° × 23.2° in Experiments 1, 2, and 3, whereas it was 72.2° × 44.7° in Experiments 4 and 5. A chin rest (T.K.K. 930a; Takei Scientific Instruments Co., Ltd., Japan) was used to minimize head and body movements. All lights in the room were turned off, and the door of the booth was covered with a blackout curtain to eliminate visual distractions. Visual stimuli were created using MATLAB R2020b (MathWorks) and Psychtoolbox 3.0.17 and controlled using a personal computer (Elite Desk 800 G5; HP, USA) with an nVIDIA GeForce RTX 2080 graphics card.

A two-interval forced-choice (2IFC) design was used in each trial, wherein the first movie was played for 20 s, followed by the second movie, which was also played for 20 s (Figure 1). The duration of stimulus presentation was selected as 20 s to reduce the intensity of the symptoms, in accordance with our ethical rules. Longer exposure to provocative stimuli induces more symptoms of MS (Golding, 2006; Kennedy et al., 2000). The question “Which movie would be more likely to make you sick if you watched it for a long time?” was presented to the participants immediately after presenting the second movie. The participants were asked to indicate the answer by pressing one of two keys; key “1” indicated the first movie, whereas key “2” indicated the second movie. The two movies had different masking patterns that were randomly selected from the seven patterns in each trial. Thus, 21 possible pairs of masking patterns were compared twice by changing the order of their presentation, resulting in 42 trials for each observer. The order of presentation of the masking pattern pairs was counterbalanced across participants.

**Figure 1.**
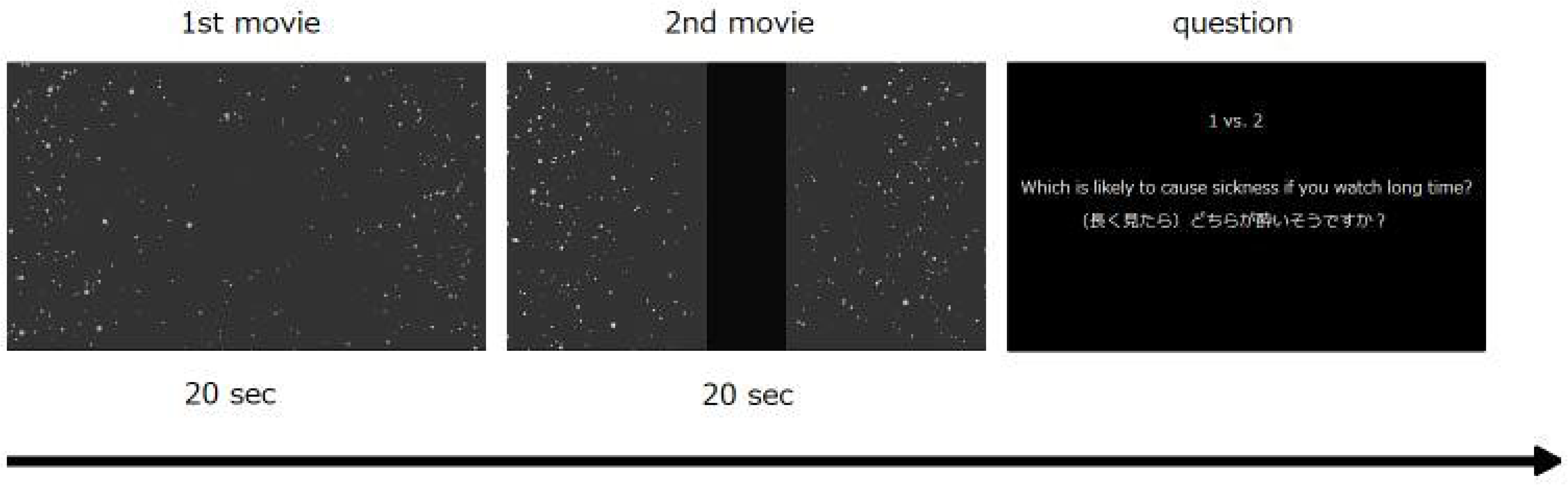
The stimuli and trial procedure. The first and second movies had different masking patterns. The question was presented in English and Japanese at the end of the second movie. The participants were instructed to answer in an unhurried manner and with care to ensure that the wrong key was not pressed.

### 2.4 Experiment 1

Experiment 1 aimed to compare different masking patterns in terms of VIMS under basic, i.e., radially expanding, optical flow. While the effects of visual screen occlusion have primarily been studied in the context of central and peripheral vision, this experiment took a different approach by systematically investigating the impact of various masking patterns. Fifty-one observers, comprising 24 males and 27 females (22.9±2.3 years), participated in Experiment 1. Thirty-three of these observers participated in Experiment1 only, whereas the remaining 18 observers participated in other experiments.

Figure 2 (a) presents the masking patterns used in Experiment 1. Pattern 1 did not involve masking. The masks were positioned at the center of the screen in Patterns 2 and 6. The width of the mask in Pattern 2 was one-sixth of the screen width, whereas the width of the mask in Pattern 6 was twice that of the width of the mask in Pattern 2. The mask was positioned at the left end of the screen in Pattern 4. In Pattern 3, the mask was positioned at the mid-point of the mask positions in Patterns 2 and 4. The width of the mask in Patterns 3 and 4 was the same as that in Pattern 2, i.e., one-sixth of the screen width. In Pattern 7, the mask was centered at the left edge of the mask in Pattern 2, and its width was twice the width of the mask in Pattern 2, i.e., one-third of the screen width, which is the same as that in Pattern 6. In Pattern 5, two masks were positioned adjacent to both the right and left edges of the mask in Pattern 6, and their widths were half of the width of the mask in Pattern 2, i.e., one-twelfth of the screen width. These masking patterns were used to evaluate the effects of the following factors on VIMS: the horizontal position of the mask, i.e., the effect of shielding the FOE or screen center; the mask width; and the number of subdivisions of the screen. In Experiment 1, a moving 3D random pattern of dots was created to be perceived as moving in the depth direction (radially expanding optical flow) at an angular velocity of 72 pixels/s along the direction of motion.

**Figure 2.**
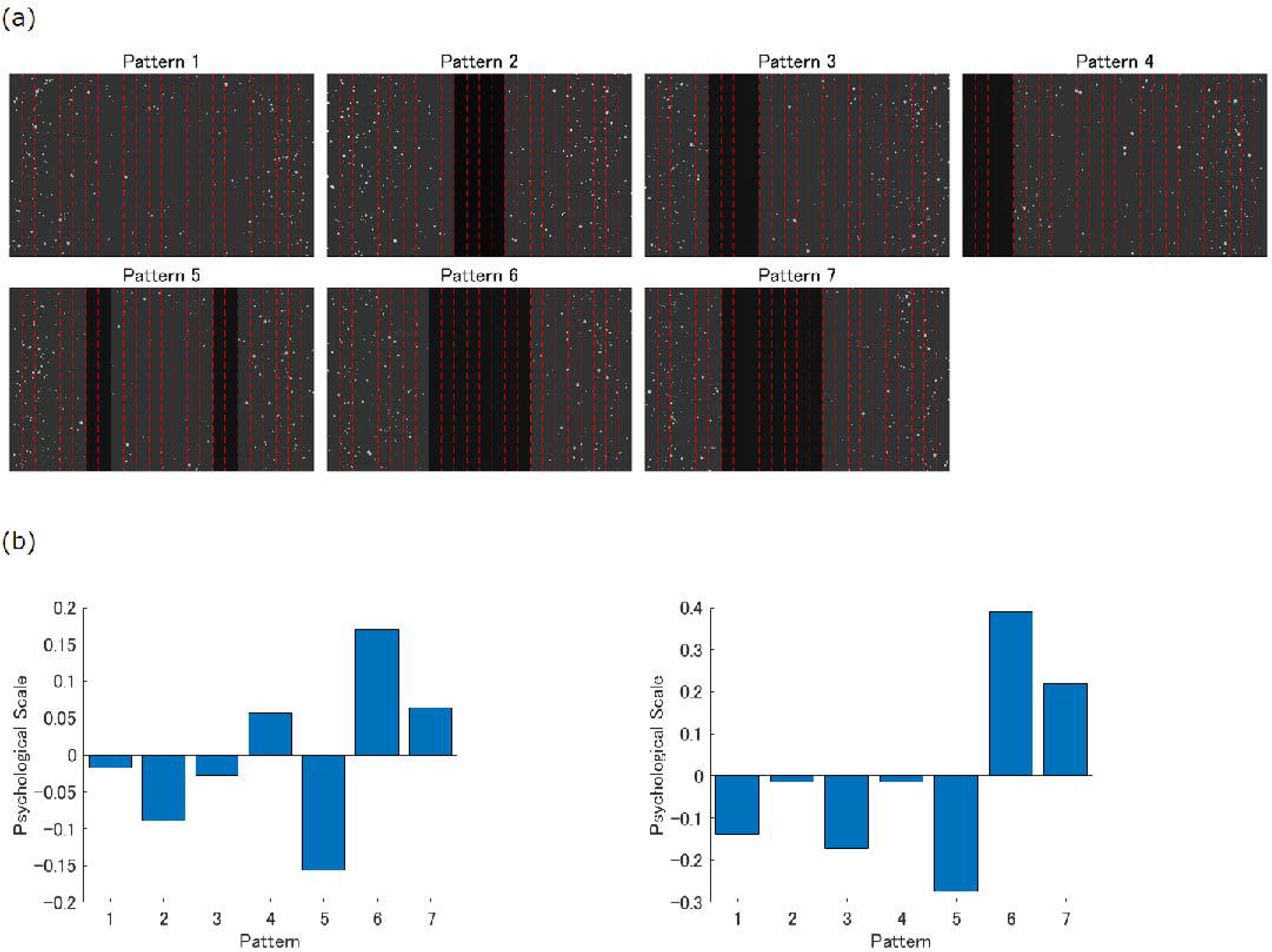
(a) Masking patterns used in Experiments 1 and 2. The masks were drawn as black rectangles. The white dots were moved to create an optical flow. The red dashed lines were not presented in the movies but are drawn in the figure to illustrate the position and width of the masks. The distance between the adjacent dashed lines was 1/24th of the screen width. (b) Results of Experiments 1 (left) and 2 (right). Psychological scales are plotted against the masking patterns. The lower and higher scales indicate that the corresponding masking pattern was likely and unlikely to induce VIMS, respectively.

### 2.5 Experiment 2

Experiment 2 aimed to investigate the effect of the masks in a situation wherein the participants were exposed to the optical flow, which is considered to be more likely to induce MS. Fifty observers, comprising 35 males and 15 females (22.8±2.1 years), participated in Experiment 2. Twenty of these observers participated in Experiment 2 only, whereas the remaining 30 observers participated in other experiments.

Experiment 2 varied from Experiment 1 in terms of the movement of random dots. The masking patterns used in Experiment 2 were identical to those used in Experiment 1 (Figure 2 (a)). The rotational movement (roll, pitch, and yaw) of a scene contributed to increased SSQ scores (Lo & So, 2001). Therefore, yawing and radial expansion movements were combined to make the stimulus more provocative compared with that of Experiment 1. The yawing motion was introduced by sinusoidally changing the parameters of the horizontal shooting direction of the camera. The amplitudes of the yawing motion were randomly jittered between 0.3 and 0.6 at a temporal frequency of 1.0 Hz to prevent the rotational motion effect from being canceled out when an observer could fully predict the motion trajectories.

### 2.6 Experiment 3

Experiment 3 aimed to investigate the effect of the number of screen divisions on VIMS. Fifty-one observers, comprising 33 males and 18 females (22.8±2.3 years), participated in Experiment 3; however, the data for one observer were excluded as the experiment was terminated halfway due to some stimulus presentation PC-related problems. Twenty-seven of these observers participated in Experiment 3 only, whereas the remaining 24 observers, including the observer whose data were removed, participated in other experiments.

The masking patterns used in Experiment 3 are illustrated in Figure 3 (a). Patterns 1 and 5 were the same as those used in Experiments 1 and 2. In Pattern 6, the two masks were located at a position beyond twice the width of the mask compared with that of Pattern 5. The width of each mask was the same for Patterns 5 and 6, i.e., one-twelfth of the screen width. The masks used in Pattern 2 were a combination of those used in Patterns 5 and 6. Pattern 7 was created by removing the second mask on the right side in Pattern 2. In Pattern 3, the centers of the two masks were located in the same position as those of the two masks in Pattern 5; however, the width of each mask was twice the width of the mask used in Pattern 5, i.e., one-sixth of the screen width. In Pattern 4, the centers of the two masks were located in the same position as those of the two masks in Pattern 5; however, the width of the masks was one-fourth of the width of the masks in Pattern 5, i.e., 1/48th of the screen width. Similar to Experiment 1, a radially expanding optical flow was used in Experiment 3.

**Figure 3.**
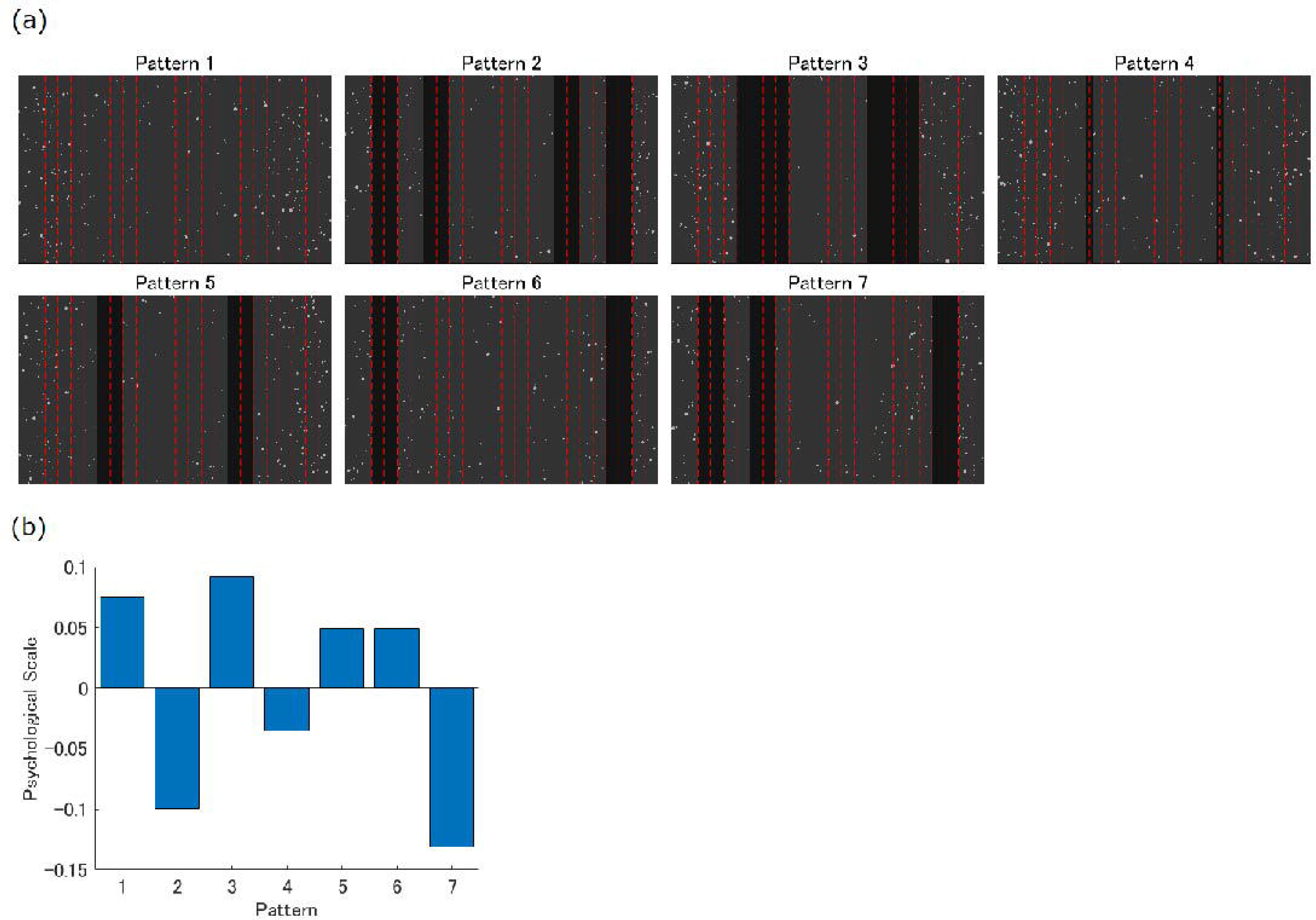
(a) Masking patterns used in Experiment 3. The masks were drawn as black rectangles. The white dots were moved to create an optical flow. The red dashed lines were not presented in the movies but are drawn in the figure to illustrate the position and width of the masks. The distance between the adjacent dashed lines was 1/24th of the screen width. (b) Results of Experiment 3. Psychological scales are plotted against masking patterns. The lower and higher scales indicate that the corresponding masking pattern was likely and unlikely to induce VIMS, respectively.

### 2.7 Experiment 4

A larger field of view has the potential to induce stronger symptoms (Van Emmerik et al., 2011). Therefore, Experiment 4 aimed to investigate the effects of masks using more intense stimuli by shortening the viewing distance. Sixty-one observers, comprising 40 males and 21 females (22.9±2.6 years), participated in Experiment 4. Thirty-three of these observers participated in Experiment 4 only, whereas the remaining 28 observers also participated in other experiments.

The masking patterns used in Experiment 4 are illustrated in Figure 4 (a). The mask was positioned at the center of the screen in Patterns 2 and 6; however, the width of the mask in Pattern 2, i.e., one-twelfth of the screen width, was half the width of the mask in Pattern 6. The width of the mask in Pattern 3 was the same as the width of the mask in Pattern 2; however, the mask in Pattern 3 was shifted by 5/48th of the width of the screen to the left. The mask on the left side in Pattern 4 was in the same position as the mask in Pattern 3, and the mask on the right side in Pattern 4 was placed symmetrically. The masks in Patterns 4 and 5 were in the same positions; however, the width in Pattern 4, i.e., 1/48th of the screen width, was half of the width of the masks in Pattern 5. Pattern 7 had four masks, and the two inner masks were in the same position as the two masks in Pattern 5.

**Figure 4.**
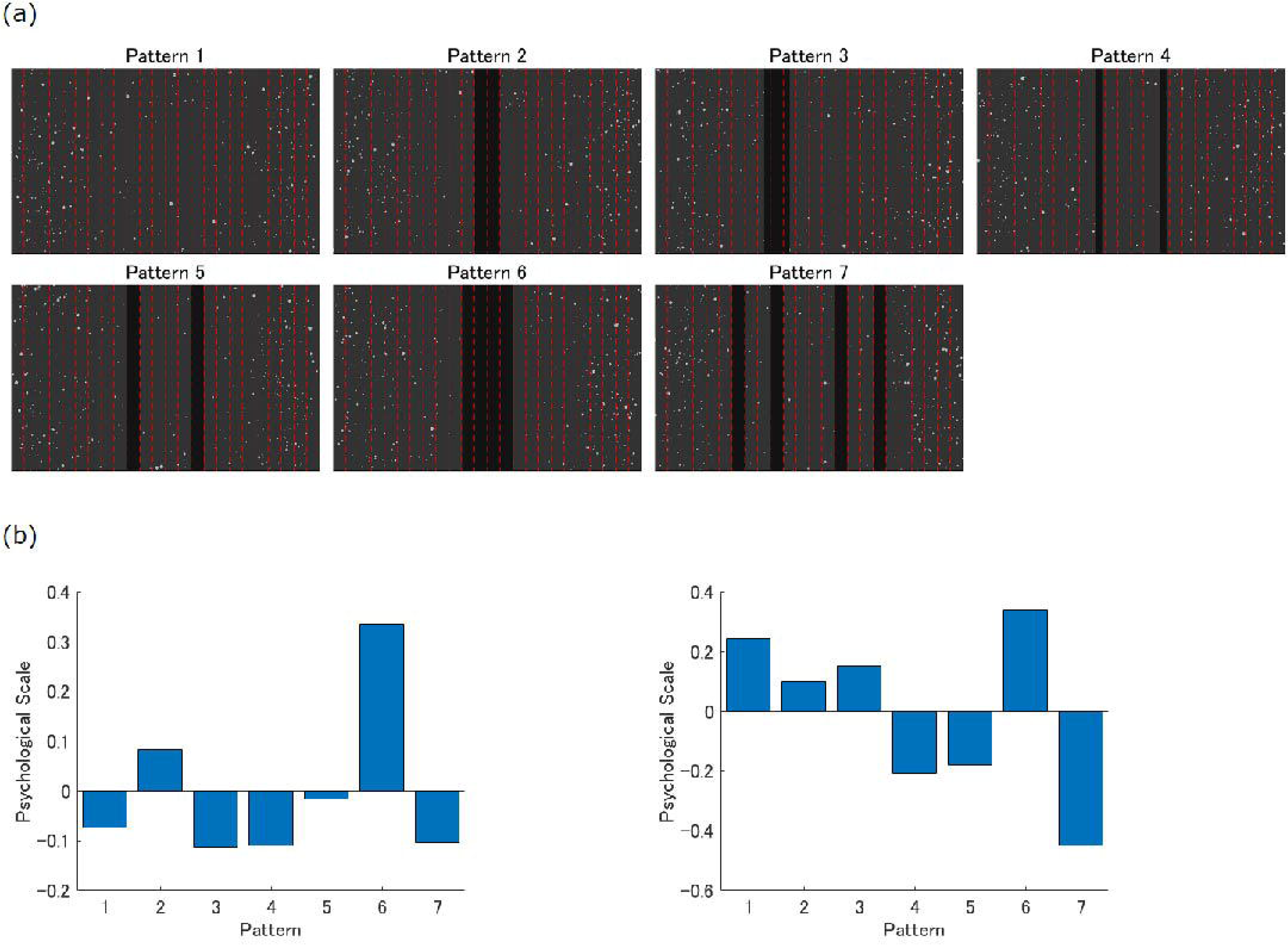
(a) Masking patterns used in Experiments 4 and 5. The masks are drawn as black rectangles. The white dots moved to create an optical flow. The red broken lines were not presented in the movies but are drawn in the figure to illustrate the position and width of the masks. The distance between the neighboring broken lines was 1/24th of the screen width. (b) Results of Experiments 4 (left) and 5 (right). Psychological scales are plotted against masking patterns. The lower and higher scales indicate that the corresponding masking pattern was likely and unlikely to induce VIMS, respectively.

Patterns 1 and 5 are the counterparts of those in Experiments 1, 2, and 3; the widths and positions of the masks in Pattern 5 were the same in terms of the visual angle. Similar to Experiments 1 and 3, a radially expanding optical flow was used in Experiment 4.

### 2.8 Experiment 5

Experiment 5 aimed to investigate the effects of masking patterns when a new optical flow was used without an FOE. Fifty observers, comprising 36 males and 14 females (22.8±2.4 years), participated in Experiment 5. Twenty-two of these observers participated in Experiment 5 only, whereas the remaining 28 observers participated in other experiments.

The masking patterns and viewing distances used in Experiment 5 were the same as those used in Experiment 4 (see Figure 4 (a)). However, the movement of random dots was varied in Experiment 4.

A yawing motion that did not include the FOE was used in this experiment. The yawing motion was introduced by sinusoidally changing the parameters of the horizontal shooting direction of the camera. The amplitude of the yawing motion was randomly jittered between 1.0 and 2.0 at a temporal frequency of 0.5 Hz to avoid any prediction or adaptation effects on the VIMS impressions.

### 2.9 Analysis

The following analysis pipeline was applied to each experiment based on Thurstone’s paired comparison method. A frequency matrix was obtained from the results of the choices of the observers for each possible pair of seven different masking patterns. The frequency matrix was converted into a percentage preference matrix by dividing the data by the number of samples. Subsequently, the percentage matrix was transformed into a z-score matrix, and the z-scores were averaged for each column. The average z-score served as a psychological scale for each masking pattern. The internal consistency of the scale value was tested to verify the mono-dimensionality of the psychological scale.

The masking patterns in each experiment had their own psychological scales. However, the differences in the masking patterns were considered in the following comprehensive analyses, and the differences in the experimental types (1-5) were ignored. First, one-way ANOVA and additional post-hoc Tukey’s honest significant difference (HSD) test were performed to test for the main effect of the number of screen subdivisions on the psychological scales. All masking patterns could be categorized into five types according to the number of screen subdivisions. For instance, Pattern 2 in Experiment 3 and Pattern 7 in Experiments 4 and 5 divided the screen into five sections (Figures 3 and 4). Thus, these three masking patterns can be classified into the same type. Second, a two-sample t-test was performed to evaluate whether the susceptibility to VIMS, i.e., the psychological scale, could be modulated when the center of the screen was occluded. In Experiments 1-4, the center of the screen matched the FOE; thus, whether the visibility of the central visual field or the FOE was essential for motion sickness was examined. All masking patterns could be categorized into two types depending on whether one of the masks occluded the center of the screen. For instance, in Experiments 1 and 2, the center of the screen was occluded in Patterns 2, 6, and 7 owing to the presence of the mask, whereas it was not occluded in Patterns 1, 3, 4, and 5 (Figure 2). Thus, Patterns 2, 6, and 7 can be classified into one type, and Patterns 1, 3, 4, and 5 can be classified into the other type. Third, a paired t-test was performed to assess whether the psychological scales were affected by the width of the masks when the positions of the masks are identical. Some thin-thick pairs were used in this test. For instance, the mask positions were identical; however, the width of the masks was different for Patterns 2 and 6 and Patterns 4 and 5 in Experiments 4 and 5 (Figure 4).

## 3. Results

### 3.1 Experiment 1

The results demonstrated that susceptibility to VIMS varied depending on how the visual field was masked or segmented, even when the optical flow remained constant. The left panel of Figure 2 (b) shows the psychological scale values for the masking patterns in Experiment 1. Statistical tests revealed that the internal consistency, i.e., the mono-dimensionality of the psychological scale, was maintained (chi-square test, χ² = 6.7104, p > 0.05). Pattern 5, which divided the visual field into three sections, had the lowest scale value, indicating that it was the most likely pattern to induce VIMS. In contrast, Pattern 6, which shielded the FOE or the center of the screen, had the highest scale value, indicating that it was the least likely pattern to induce VIMS. The comparison between Patterns 2 and 6 suggests that a wider mask may prevent the incidence of VIMS when the mask positions were identical.

### 3.2 Experiment 2

In Experiment 2, a visual motion pattern more likely to induce VIMS was employed, featuring radial expansion combined with yawing movements with some motion jitters to prevent prediction and adaptation effects on VIMS. The results of Experiment 2 were similar to those of Experiment 1. The right panel of Figure 2 (b) shows the psychological scale values for the masking patterns in Experiment 2. Statistical tests confirmed the internal consistency (chi-square test, χ² = 13.6413, p > 0.05). Similar to the results of Experiment 1, Pattern 5 was found to be the pattern most likely to induce VIMS, whereas Pattern 6 was found to be the pattern least likely to induce VIMS. In addition, the comparison between Patterns 2 and 6 revealed that a wider mask prevented the incidence of VIMS in the same position.

### 3.3 Experiment 3

The number of screen divisions affected the susceptibility to VIMS, as suggested in previous experiments. Figure 3 (b) shows the psychological scale values for the masking patterns in Experiment 3. Statistical tests confirmed the internal consistency (chi-square test, χ² = 10.8785, p > 0.05). The patterns with a greater number of screen subdivisions, such as Patterns 7 and 2, were more likely to induce VIMS. The comparison among Patterns 3, 4, and 5 revealed that a wider mask prevents the incidence of VIMS when the mask positions were identical. These results are consistent with those of previous experiments.

### 3.4 Experiment 4

The findings remained consistent even when the stimuli were amplified by increasing the screen size and enhancing the impression of motion. The left panel of Figure 4 (b) shows the psychological scale values for the masking patterns in Experiment 4. Statistical tests confirmed the internal consistency (chi-square test, χ² = 18.0350, p > 0.05). As observed in the previous experiments, (1) masking patterns that occluded the FOE or the center of the screen, such as Patterns 6 and 2, were less likely to induce VIMS, and (2) the comparisons between Patterns 4 and 5 and between Patterns 2 and 6 revealed that wider masks prevented the incidence of VIMS when the mask positions were identical. However, the masking pattern with more extensive screen divisions was not found to be more likely to induce VIMS. The psychological scale of the masking pattern with the highest number of screen divisions (Pattern 7) was low, but not the lowest.

### 3.5 Experiment 5

The findings remained consistent even when a new optical flow without FOE was employed. The right panel of Figure 4 (b) shows the psychological scale values for the masking patterns in Experiment 5. Although the statistical test did not confirm the internal consistency (chi-square test, χ² = 29.3414, p < 0.05), the following trends were observed. First, the observers were found to be more susceptible to VIMS when the masks divided the screen into a greater number of sections (Patterns 7, 4, and 5). Second, the masking pattern that shielded a wide area around the center of the screen, i.e., Pattern 6, was found to be the pattern least likely to induce VIMS. Third, the comparisons between Patterns 4 and 5 and between Patterns 2 and 6 revealed that wider masks prevented the incidence of VIMS when the mask positions were identical. These trends were consistent with the results of the previous experiments.

### 3.6 Results of the comprehensive analysis

This section reviews and summarizes all five experiments and provides general insights into the relationship between the susceptibility to VIMS and the types of optical flow and mask patterns (visual field appearances). The results of each experiment are summarized in Figure 5. First, the participants were more susceptible to VIMS when the screen was divided into a greater number of sections by masks. Figure 5(b) shows that the masking patterns with more reddish markers, indicating that the masks divided the screen into more sections, had lower psychological scales.

**Figure 5.**
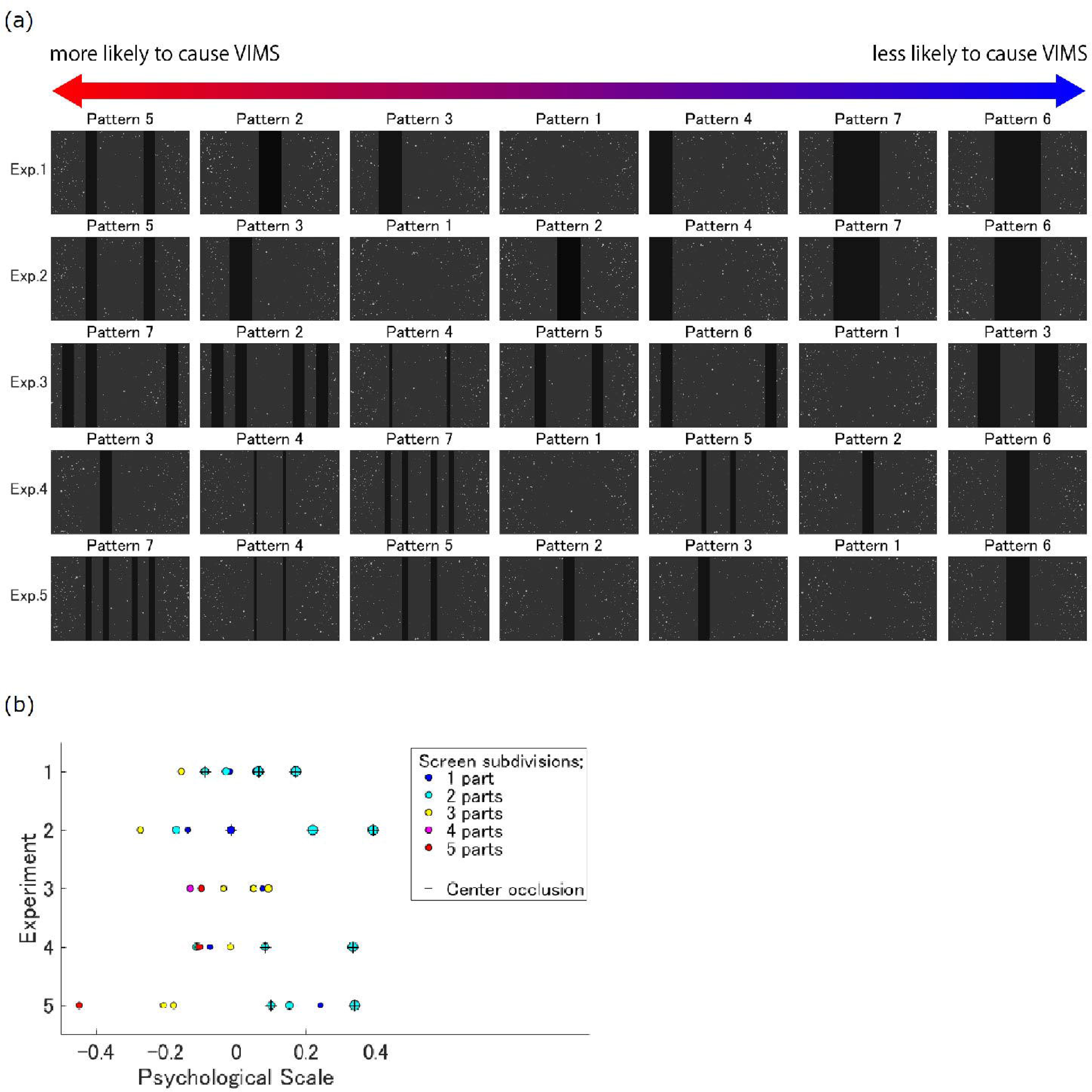
Summary of all experiments. (a) Masking patterns were sorted according to their psychological scales for each experiment. The masks with high psychological scales are less likely to induce VIMS, whereas the masks with low psychological scales are more likely to induce VIMS. (b) For each experiment, each masking pattern was plotted according to its psychological scale. More reddish markers indicate that the masks divided the screen into more sections (red, 5 sections; magenta, 4 sections; yellow, 3 sections; cyan, 2 sections; blue, one section). The size of the marker reflects the width of the mask. The ‘+’ sign is overwritten for the masking patterns where one of the masks shielded the center of the screen.

Second, the participants were found to be less susceptible to VIMS when the masks were wider when the mask positions were identical. Third, the participants were found to be less susceptible to MS when the center of the screen was masked. Figure 5(b) shows that the masking patterns with “+” markers had higher psychological scales. Statistical analyses were performed to validate these findings.

One-way ANOVA revealed that the psychological scales differed significantly depending on the number of screen subdivisions (F_(4,30)_ = 3.89, p < 0.05). Post-hoc Tukey’s HSD test revealed that the psychological scales of the masking patterns that divided the screen into two sections, such as Pattern 2 in Experiment 4, were significantly higher than those of the masking patterns that divided the screen into five sections, such as Pattern 7 in Experiment 4 (p < 0.05). Although a significant difference was observed only between these pairs, the mean psychological scales decreased with the number of divided screens when the screen was divided by masks (Figure 6(a)).

**Figure 6.**
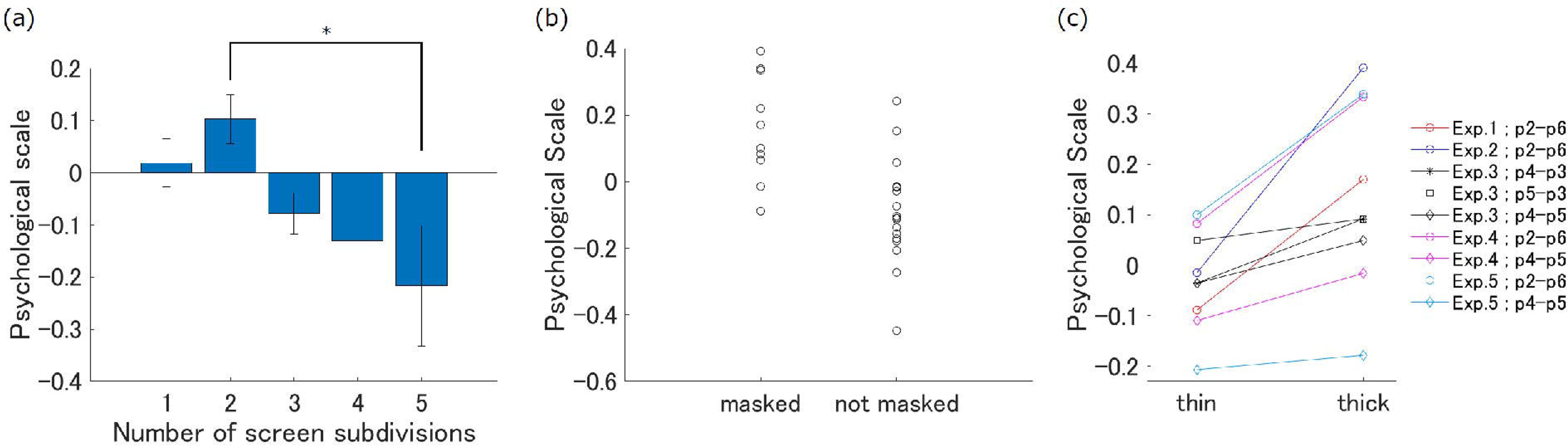
Psychological scales were combined across the experiments to investigate the factors affecting the psychological scales. (a) The mean psychological scales for different numbers of screen divisions. The error bars represent the standard deviation of the mean. *p < 0.05. (b) Psychological scales for the masking patterns in which the center of the screen was occluded by the mask and those in which the center of the screen was not occluded by the mask. Each circle corresponds to each masking pattern of a given experiment. (c) Psychological scales for pairs of masking patterns in which the positions of the masks were identical. Each circle corresponds to each masking pattern of a given experiment.

The two-sample t-test revealed that masking patterns that occluded the center of the screen had significantly higher psychological scales than those that did not (p < 0.0005), with a mean psychological scale of 0.1598 and -0.0888 for masking patterns that did and did not occlude the center of the screen, respectively (Figure 6(b)).

The paired-sample t-test revealed that the psychological scales for thick masks were significantly higher than those for thin masks when the mask positions were identical (p < 0.005). This trend was observed in all pairs (Figure 6(c)).

## 4. Discussion

### 4.1 Discussion of findings

VIMS is an urgent issue that must be addressed for the widespread adoption of emerging technologies such as virtual/augmented reality (VR/AR), and for developing comfortable autonomous vehicles. Previous studies have extensively focused on identifying the types and characteristics of motions that are more likely to induce MS. In contrast, less attention has been paid to how contextual factors, such as framing, visual field segmentations, occlusions, or the appearance of graphical user interface (GUI) windows, may change the feeling of MS even under identical motion patterns. To bridge this gap, the present study investigated the contextual effects of visual field segmentations and occlusions on MS susceptibility through five experiments.

Our paired-comparison-based analyses revealed the followings: (1) the psychological scales decreased, i.e. the likelihood of inducing VIMS increased, as the number of screen divisions by masks, (2) the masking patterns that occluded the center of the screen had significantly higher psychological scales, i.e. a lower possibility of inducing VIMS, compared to those that did not, and (3) the psychological scales of thick masks were higher than those of thin masks when the mask positions were identical.

The first finding suggests that VIMS is more likely to occur when the screen is divided into a greater number of sections by the masks, which leads to more frequent interruption of the trajectory of each dot. However, it is unclear why the fragmentation of the visual field tends to induce VIMS. As described in the Introduction, the sensory conflict theory states that a mismatch of information between or within sensory modalities leads to MS. Mismatch, occlusion, and interruption remind us of the Poggendorff illusion, an optical illusion wherein observers perceive two-line segments as being misaligned when an obliquely oriented line is bisected by an opaque rectangular occluder or two parallel lines. The kinetic Poggendorff illusion (Nihei, 1974, 1976; Wenderoth & Johnson, 1983;

Fineman & Melingonis, 1977; Watamaniuk, 2005) is the dynamic version of the Poggendorff illusion. When a dot moves obliquely behind vertical parallels or an opaque rectangle, the trajectories before and after the intersection are perceived as being misaligned. According to Nihei (1974), the magnitude of the misalignment is influenced by the angle of the oblique motion from the vertical direction and is a function of the motion velocity. In the present study, many dots showed oblique radial motion from the focal point at the center of the screen in different directions behind the rectangular masks, which could lead to various perceived misalignments, i.e. unstable zig-zag dot movements. These misalignments can induce VIMS, as predicted by the sensory conflict theory.

However, in the present study, the velocity of the dot motion was relatively slower than that in previous studies; therefore, the magnitude of the illusion, if any, was very small. Furthermore, although the kinetic Poggendorff illusion has been reported to be observed in two dimensions, it is unknown whether such an illusory misalignment would be observed in 3D in the present study. In addition, we provide a counterexample of the illusory misalignment hypothesis. According to Talasli and Inan (2015), misalignment increases with the width of the occluder in the static Poggendorff illusion. If this is also applicable to moving dots in a 3D space, observers are more likely to experience VIMS when the width of the mask is thicker. However, contrary to this prediction, the findings of the present study indicated that thicker masks had higher psychological scales, indicating that they are less likely to induce VIMS when the mask positions are identical (Figure 6(c)). Thus, it is difficult to explain the first finding of the present study based on the illusory misalignment hypothesis, and the mechanisms underlying these findings remain unclear.

The second finding suggests that the masking patterns that occluded the center of the screen were less likely to induce VIMS than those that did not. This was observed in Experiments 1 and 4, where the FOE corresponded to the center of the screen, and in Experiment 5, where the FOE was not present, suggesting that the visibility of the center of the screen is the factor inducing VIMS, not the visibility of the FOE. This finding seems to be inconsistent with the findings of previous studies that investigated the relationship between VIMS-related phenomena and central/peripheral vision. Kim and Kim (2019) compared the SSQ scores for peripheral vision (the central 10° view of each eye was masked), central vision (the view of each eye was masked except for the central 10°), and full vision.

They reported that the SSQ scores were the highest (indicating most severe symptoms of VIMS) for peripheral vision and the lowest (indicating least severe symptoms) for central vision. In the present study, the width of the central mask was about 6.7° of the visual angle for Pattern 2 and about 13.4° of the visual angle for Pattern 6 in Experiments 1 and 2. These numerical values are comparable with the peripheral vision conditions in the study by Kim. These contradictory results may be a result of the differences in masked parts of the visual field as well as the differences in methods used for assessing the VIMS symptoms and stimulus presentation (Kim and Kim (2019) used head-mounted displays). The mask used in the present study, i.e., a vertical rectangle, occluded the central field and the upper and lower portions of the center of the screen, including the peripheral field. Additional masking of these areas may reduce the possibility of inducing VIMS. As shown in the Supplementary Information, horizontal eye positions had higher variability and the frequencies of optokinetic nystagmus (OKN) and horizontal saccades were higher when the area around the center of the screen was vertically masked. Given that the psychological scales were higher in such situations, VIMS symptoms cannot be attributed to the increase in the eye movements. The Eye Movement theory links VIMS with OKN (Ebenholtz et al., 1994) and previous studies have shown that the severity of VIMS is correlated with the frequency (Hu & Stern, 1998) and slow phase velocity (Ji et al., 2009) of OKN. However, a more recent study (Nooij et al., 2017) that used more advanced experiments and analyses demonstrated that these parameters did not contribute significantly to VIMS. In any case, the reason why vertical masking of the center of the screen reduced the possibility of inducing VIMS in our experiments cannot be explained using eye movements. However, the finding that masking patterns that vertically shield the area around the center of the screen are less likely to induce sickness is novel and important in that it can aid in designing visual displays, dynamic advertisements, child-friendly movies, and motion-sickness-free web pages.

The third finding indicates that thin masks are more likely to induce VIMS than thick masks when the mask positions are identical. As described previously, this result is inconsistent with the illusory misalignment hypothesis. A simple explanation for this could be that more moving dots would enhance the impression of VIMS when thinner masks occluded only limited parts of the visual field. However, if a larger number of visible dots were the primary factor in inducing VIMS, the no-masking condition (Pattern 1 in all experiments), which presented the maximum number of visible dots, would have been expected to induce the highest VIMS susceptibility. However, this was not the case. Instead, the results suggest that the segmentation of the visual field had a stronger effect on VIMS induction than the number of visible dots. Or more precisely, the findings indicate that no single factor, such as the number of visual field divisions or visible dots alone, can fully explain the observed results. Rather, some interactions between visual field segmentations and the number of visible dots may play a critical role in shaping the perception of VIMS. Future follow-up studies are required to explore these interactions and provide a more comprehensive understanding of the mechanisms underlying VIMS induction.

### 4.2 Limitations and future research

This study examined the impact of visual field masking and fragmentation on susceptibility to VIMS by manipulating visual modality-related parameters, such as masking patterns and optical flow patterns. However, the effect of visual field masking and fragmentation on the susceptibility to MS in the presence of other motion information from the vestibular and somatosensory systems is unclear. For instance, in a situation wherein an individual is seated in the rear seat of a moving car, the view must be occluded or divided by the front seats and pillars. Such visual occlusion will affect the visual information regarding motion without affecting information from other sensory modalities. Thus, the discrepancy between the visual and other sensory modalities should be investigated further. Since the sensory conflict theory states that a mismatch between visual, vestibular, and somatosensory information leads to the incidence of MS, the masking and fragmentation of the visual field should affect susceptibility to MS. This should be addressed in future studies.

This study has some limitations. The participants were instructed to indicate which of the two movies (20 s each) was more likely to induce sickness in the paired comparison experiments (Please note that this setup was ethically required for wellness of participants by preventing them from real sickness). Owing to the forced-choice nature, the participants were instructed to answer even if they did not experience VIMS. The short exposure to the optical flow stimulus in the experiments may not have had a true impact on the induction of VIMS. Therefore, the present results may be criticized as they do not necessarily guarantee a relationship between masking patterns and real VIMS. Nevertheless, the findings of the present study seem reliable as the same results were consistently observed over different experiments using different motions (e.g., expanding and/or yawing) and across observers. It would be beneficial to investigate whether the present findings can be reproduced using a VIMS severity measure, such as the SSQ, to improve the robustness of these findings.

### 4.3 Conclusion

In conclusion, the results indicate that (1) VIMS is more likely to occur when the screen is divided into a greater number of sections by the masks, (2) the masking patterns that occluded the center of the screen were less likely to induce VIMS than those that did not, and (3) thin masks are more likely to induce VIMS than thick masks when the mask positions are identical. These findings provide valuable insights for a wide range of industrial applications, including the development of safer VR/AR environments, no-dizziness GUI windows, and user-friendly movie viewing systems, as well as the design of more comfortable automotive systems, with minimizing VIMS.

## Author contributions

**Takatsugu Aihara**: Conceptualization, Methodology, Software, Investigation, Data curation, Formal analysis, Visualization, Writing-original draft, Writing-review and editing; **Sachiko Yamada**: Conceptualization, Methodology; **Hiroshi Ban**: Conceptualization, Methodology, Software, Writing-review and editing, Funding acquisition, Supervision, Project administration.

## Data availability

Data will be made available on request.

## Acknowledgments

This study was supported by the Honda-NICT collaborative research Fund (TA, SY, HB) and JSPS Kakenhi (21H00968, 21K18572) to HB. The funders had no role in the study design, data collection, and analysis, decision to publish, or manuscript preparation.

DeepL PRO and ChatGPT-3.5 services were used in part for English proofreading. The authors took special care to ensure that no changes were made to the original content by adding a clear instruction such as "…, keeping the meanings the same" to the chatbot’s prompt, and by double-checking the updated manuscript.

We thank Miho Hayashi for assistance with data acquisition and Naomi Murakami for organizing the experimental schedule by contacting the participants.

## Appendix A. Supplementary information

Supplementary analyses and results are included in the Supplementary Information. Movie files depicting examples of the visual stimuli are also included in the Supplementary Information. For example, “Exp1_pat1_L.avi” is the movie for the masking pattern 1 in Experiment1, which was shown to participant’s left eye.

