## Supplementary Information for "The Effect of Visual Field Occlusion on Visually Induced Motion Sickness"

The Supplementary Information presents the results of the analyses of the eye position data recorded in Experiment 1. These analyses aimed to assess whether the masking patterns affect eye movements and clarify whether visually induced motion sickness (VIMS) is linked to eye movements.

According to the Eye Movement theory (Ebenholtz et al., 1994), optokinetic nystagmus (OKN), a sequence of "slow" pursuit and "fast" backward movements (saccades) typically observed when viewing a horizontally moving grating pattern, is a contributing factor to VIMS. Previous studies have demonstrated that the severity of VIMS is positively correlated with the frequency (Hu & Stern, 1998) and slow phase velocity (Ji et al., 2009) of OKN; however, some studies have also reported that these parameters are not significant predictors of VIMS (Nooij et al., 2017). The incidence of OKN in the present study is unlikely as expanding optical flow stimuli rather than horizontally rotating stripe stimuli were used in the present study. The number of OKNs in Experiment 1 was low; nevertheless, the OKN parameters were compared between the masking patterns.

In the next step of the eye movement analysis, the mean and standard deviation (SD) of the eye position distribution were compared between the masking patterns. Finally, the horizontal saccades were compared across masking patterns as the distribution analysis and visual inspection of the temporal changes in eye position (not shown) suggested that the participants moved their eyes from side to side to avoid the mask when the mask covered the center of the screen.

### MATERIALS AND METHODS

In Experiment 1, the eye position data were recorded for all 51 participants using EyeLink II (SR Research Ltd., Ottawa, Canada) while watching the movie. The data were sampled at 2000 Hz and processed offline using MATLAB.

Data pre-processing was performed as follows. The pupil was occluded during blinking; therefore, not a number (NaN) was allocated to the eye position data on MATLAB. Thus, the timing of the blinks was determined to identify the NaN. The data from the 75-ms period immediately before and after blinking were removed from the later analysis, i.e., data from those periods were replaced by NaN, to avoid contamination of the blink-related changes in eye positions. The participants with NaN in $\geq$ 70 % of the samples were excluded from the analysis after the pre-processing step. Among the 51 participants, 35 were analyzed and 16 were discarded.

#### 1. OKN parameters

OKN analysis was performed as described in the study by Behrens and Weiss (1992). The angular positions of the eyes were first low-pass filtered (finite impulse response (FIR) filters with a Hamming window) at a cut-off frequency of 25 Hz. The filtered data were subsequently downsampled to 250 Hz. The eye velocity and acceleration were calculated from the pre-processed position data as the first and second derivatives, respectively.

The OKN slow phase was defined as follows. In the first step, saccade onsets and offsets were identified as an OKN slow phase is observed between the end of a saccade and the beginning of the subsequent saccade. The beginning of a saccade was determined by the first traverse of a given acceleration threshold (1200 deg/s in the present case) when the absolute values of the acceleration $|A_e|$ in the succeeding time interval $\Delta T_1$ (12 ms, corresponding to the three samples in the present study) were above the threshold. The end of a saccade was determined by the traverse below the threshold when $|A_e|$ in the succeeding time interval $\Delta T_2$ (16 ms, corresponding to the four samples in the present study) was sub-threshold. In the second step, each interval between consecutive saccades was determined to meet the requirements of the OKN slow phase. The pursuit eye movements during the OKN slow phase must extend radially outward from the center of the screen as expanding optical flow stimuli were used in the present study. The requirements for satisfying this condition are as follows: (1) The angle formed by the line connecting the viewing point to the center of the screen against a horizontal line must remain nearly constant during the slow phase period. (2) The distance from the center of the screen must be increased monotonically during the slow phase period. The OKN frequency, which indicated the number of OKNs present in 1 s, was calculated after identifying the OKN slow phases using the abovementioned procedure. In addition, the OKN slow phase velocity (SPV), which indicated the mean velocity during each slow phase, was calculated.

One-way analysis of variance (ANOVA) was performed to determine the main effect of the masking patterns on the OKN frequency and SPV to investigate the statistical effect of the masking patterns on the two OKN parameters. Tukey's honest significant difference (HSD) test was performed if significant differences were observed.

**2. Distribution of eye position**

Bivariate histograms were created to visually investigate the changes in the eye position with masking patterns using the eye position data, which comprised horizontal and vertical positions.

Statistical tests were performed separately for the horizontal and vertical directions. The mean and SD were calculated for the measures of the horizontal and vertical distributions of right and left eye positions. Differences in the parameters of the distribution between the masking patterns were tested using one-way ANOVA, followed by multiple comparisons using Tukey's HSD test.

**3. Horizontal saccade frequency**

A saccade interval was defined as an interval whose velocity, averaged over the last 20 ms, was greater than 100 deg/s. As described previously, only the case in which the participants moved their eyes from side to side to avoid the mask when the mask covered the center of the screen were considered in this study. Therefore, the target horizontal saccade was defined as a saccade whose horizontal displacement was greater than the width of the narrowest mask, i.e., the width of Pattern 5 in Experiment 1 = one-twelfth of the screen width. The horizontal saccade frequency, which indicated the number of horizontal saccades occurring in 1 s, was calculated. Differences in the horizontal saccade frequency between the masking patterns were tested using one-way ANOVA, followed by multiple comparisons using Tukey's HSD test.

**RESULTS**

**1. OKN parameters**

One-way ANOVA revealed that the OKN frequencies differed among the masking patterns (Figure S1(b), left panel). Post-hoc Tukey's HSD test revealed that the OKN frequencies of Patterns 6 and 7 were significantly higher than those of the other masking patterns, except for Pattern 2 ($p < 0.05$). As the center of the screen was shielded vertically in Patterns 2, 6, and 7 (Figure S1(a)), OKN was considered to have increased in the patterns covering the center of the screen. One-way ANOVA revealed no significant difference in the OKN SPV between the masking patterns (Figure S1(b), right panel).

(a)

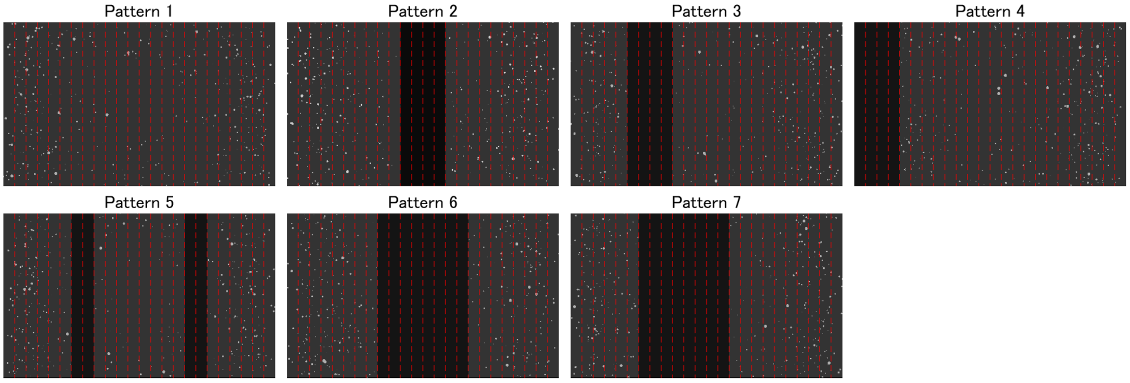


(b)

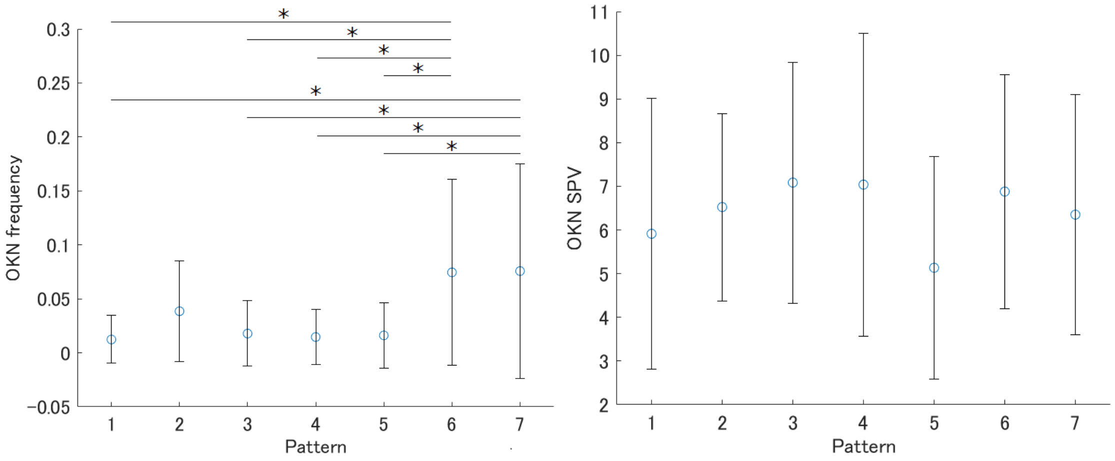


Figure S1. (a) Masking patterns in Experiment 1 (copied from Figure 2(a) in the main manuscript). (b) The frequency (times/s, left panel) and slow phase velocity (SPV) (deg/s, right panel) of optokinetic nystagmus (OKN) were plotted against the masking patterns. The error bars represent the standard deviation, and the asterisks indicate significant ($p < 0.05$) differences between the masking patterns.

### 2. Distribution of eye position

Figure S2 presents the bivariate histograms of the right eye positions. Left eye position data yielded similar results (data not shown). The histograms suggest that the observers gazed at a position slightly below the center of the screen ([1280 720]) for the majority of the time for all masking patterns. However, the high-frequency area (yellowish area in Figure S2) was spread horizontally when the center of the screen was vertically occluded (Patterns 2, 6, and 7), suggesting that the horizontal variability of the eye position was increased. In addition, the horizontally wide distributions in Patterns 2, 6, and 7 may reflect the bimodal distributions in some observers to avoid observing the masked area.

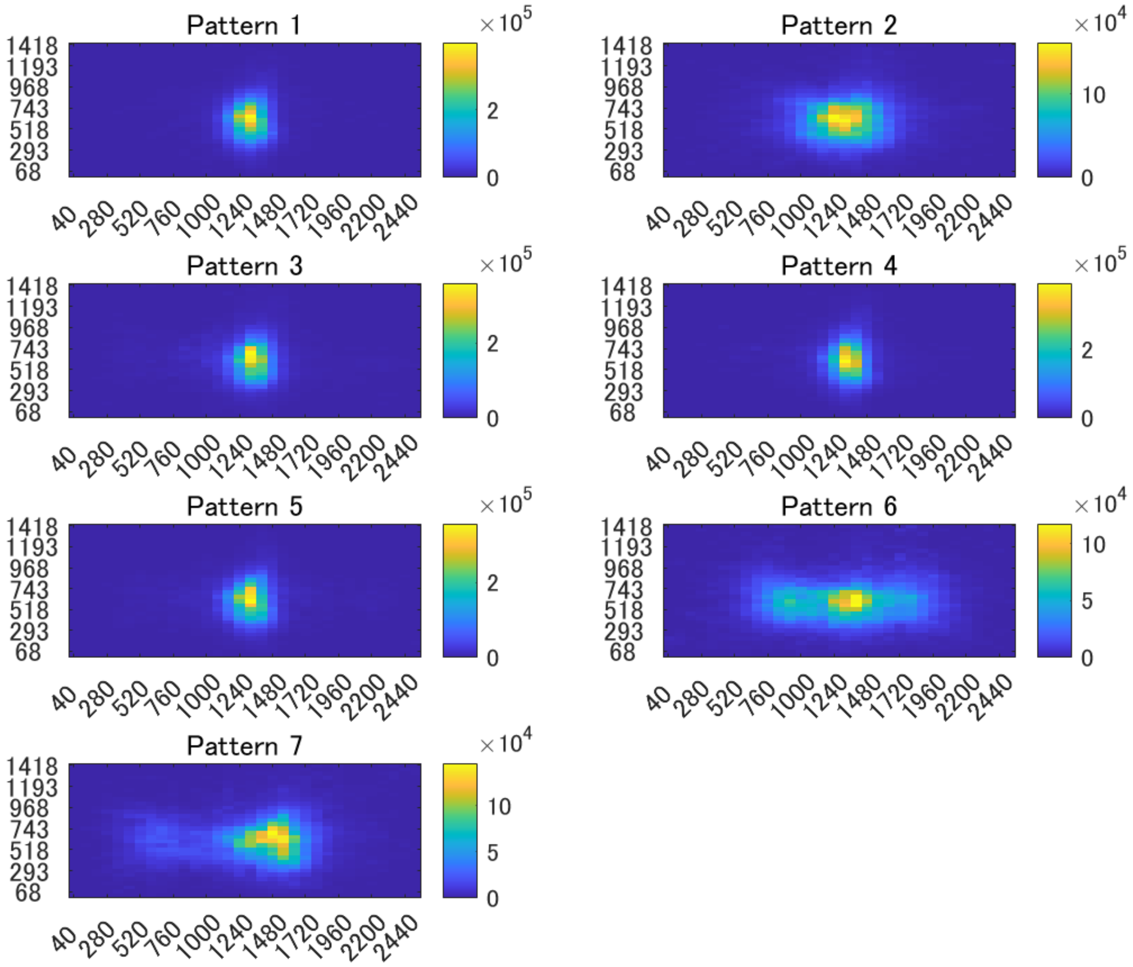


Figure S2. Bivariate histograms (summation of all thirty-five observers) of right eye positions for the seven masking patterns.

One-way ANOVA revealed that the mean horizontal eye position did not differ significantly between the masking patterns [$F_{(6,238)} = 1.15$, $p > 0.05$ for left eye; $F_{(6,238)} = 1.46$, $p > 0.05$ for right eye]. As shown in the left panel of Figure S3, the eyes of the participants were positioned around the center of the screen (x = 1280) regardless of the masking pattern. The SD of the horizontal eye position revealed a significant difference between the masking patterns [$F_{(6,238)} = 26.98$, $p < 10^{-23}$ for the left eye; $F_{(6,238)} = 28.25$, $p < 10^{-24}$ for the right eye]. Post-hoc Tukey's HSD test revealed that the SDs of Patterns 6 and 7 were significantly larger than those of the other patterns for both eyes and that the SD of Pattern 2 was significantly larger than those of Patterns 1 and 4 ($p < 0.05$; Figure S3, right panel). As shown in Figure S1(a), Patterns 2, 6, and 7 shielded the center of the screen vertically, suggesting that masking patterns that occlude the center of the screen are likely to increase the horizontal variability of the eye positions. In addition, the masking patterns with a wider central mask, such as Patterns 6 and 7, had more variability in the horizontal eye positions than the masking pattern with a narrower central mask, such as Pattern 2, suggesting that participants sometimes moved their eyes from side to side to avoid the mask when the mask covered the center of the screen.

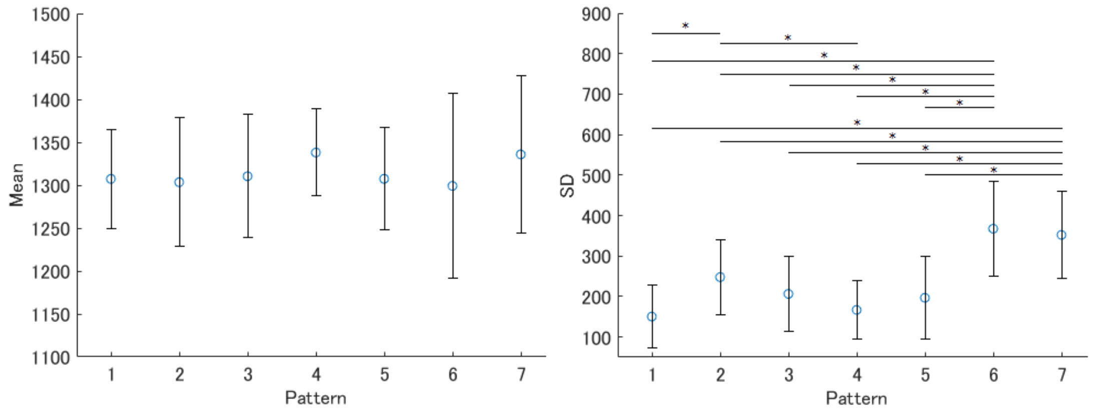


Figure S3. The mean (left panel) and standard deviation (SD; right panel) of the horizontal right eye positions (pixel position) plotted against masking patterns. The error bars represent the SDs of the participants. The asterisks indicate significant ($p < 0.05$) differences between the masking patterns. Similar results were obtained for the left eye.

One-way ANOVA revealed that the mean vertical eye position did not show a significant difference between the masking patterns [$F_{(6,238)} = 0.04$, $p > 0.05$ for the left eye; $F_{(6,238)} = 0.1$, $p > 0.05$ for the right eye]. Similarly, the SD of the vertical eye position did not differ significantly between the masking patterns [$F_{(6,238)} = 0.75$, $p > 0.05$ for the left eye; $F_{(6,238)} = 0.2$, $p > 0.05$ for the right eye]. These results suggest that the vertical eye position was not affected by the masking patterns.

#### 3. Horizontal saccade frequency

One-way ANOVA revealed that the horizontal saccade frequency differed significantly between the masking patterns [$F_{(6,112)} = 10.34$, $p < 10^{-8}$]. Post-hoc Tukey's HSD test revealed that Patterns 6 and 7 increased horizontal saccades compared with patterns other than Pattern 2. This suggests that masking patterns that occluded the center of the screen increased the horizontal saccades.

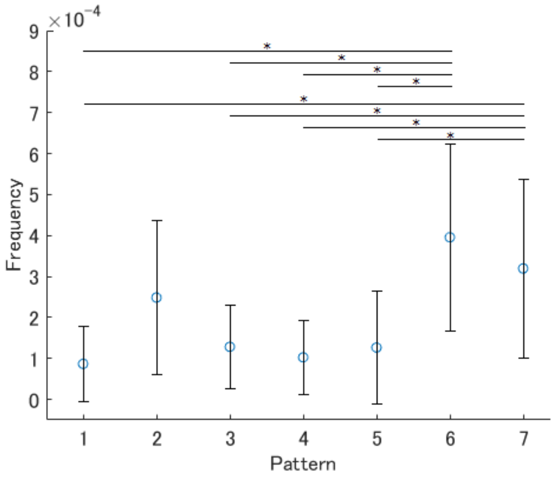


Figure S4. Horizontal saccade frequency (times/s) was plotted against masking patterns. The error bars represent the standard deviation of the participants. The asterisks indicate significant ($p < 0.05$) differences between the masking patterns.

### DISCUSSION

If eye movements, such as OKN, induce VIMS, as suggested by some previous studies, it is predicted that masking patterns, such as Patterns 6 and 7 in Experiment 1, will be more likely to induce VIMS as increased eye movements, including OKN, were observed in these patterns. However, as described in the main manuscript (for example, the left panel of Figure 2b), the behavioral results of the paired comparison in Experiment 1 were contrary to expectations. Thus, the data obtained in the present study does not support the eye movement theory of VIMS.
